# Identification of novel chemical scaffolds that inhibit the growth of *Mycobacterium tuberculosis* in macrophages

**DOI:** 10.1101/2021.09.28.462274

**Authors:** Sara Ahmed, Alyssa Manning, Lindsay Flint, Divya Awasthi, Tanya Parish

## Abstract

*Mycobacterium tuberculosis* is an important global pathogen for which new drugs are urgently required. The ability of the organism to survive and multiply within macrophages may contribute to the lengthy treatment regimen with multiple drugs that are required to cure the infection. We screened the MyriaScreen II diversity library of 10,000 compounds to identify novel inhibitors of *M. tuberculosis* growth within macrophage-like cells using high content analysis. Hits were selected which inhibited the intramacrophage growth of *M. tuberculosis* without significant cytotoxicity to infected macrophages. We selected and prioritized compound series based on their biological and physicochemical properties and the novelty of the chemotypes. We identified five chemical classes of interest and conducted limited catalog structure-activity relationship studies to determine their tractability. We tested activity against intracellular and extracellular *M. tuberculosis*, as well as cytoxicity against murine RAW264.7 and human HepG2 cells. Benzene amide ethers, thiophene carboxamides and thienopyridines were only active against intracellular bacteria, whereas the phenylthiourea series was also active against extracellular bacteria. One member of a phenyl pyrazole series was moderately active against extracellular bacteria. We identified the benzene amide ethers as an interesting series for further work. These new compound classes serve as starting points for the development of novel drugs to target intracellular *M. tuberculosis*.

## INTRODUCTION

Tuberculosis (TB) is the leading cause of death from a bacterial infection worldwide. Despite the availability of chemotherapy, TB is responsible for >10 million infections annually and >1 million deaths (1,2). The increasing incidence of drug failure for TB due to widespread drug-resistant infections underscores the urgent need for new drugs and new bacterial targets for *M. tuberculosis*.

*M. tuberculosis* is an intracellular pathogen which can survive and replicate within macrophages; this property relies on the ability to block normal bactericidal activities and turns the macrophage into a niche for replication (3–6). The physiology and metabolism of *M. tuberculosis* within macrophages is very different from extracellular bacteria and so the complement of essential genes differs (7–9). From a drug discovery perspective, this means that intracellular bacteria have additional essential processes that can be targeted by chemical inhibition.

Whole cell screening has been broadly used to identify new drug candidates for tuberculosis (10–12). In contrast to target-based drug discovery, in phenotypic screens every essential pathway is a potential drug target. This greatly increases the likelihood of finding inhibitors and overcomes the barrier of obtaining cell-penetrant molecules, which is often a limitation of biochemical screens (13). However, conditionally essential genes that are required for *M. tuberculosis* growth within macrophages or *in vivo*, may be overlooked in phenotypic screens against axenically-cultured *M. tuberculosis*. High-throughput screening in conditions that closely resemble the intracellular environment have the potential to uncover novel antimicrobials that target these conditionally essential genes, in addition to discovering compounds targeting the host cells to increase bacterial elimination (14,15).

High-content imaging screening is a powerful technology to identify active compounds by monitoring both bacterial and macrophage cell numbers simultaneously and in the same wells, thus leading to more reliable data and a quicker assessment of compound attractiveness. We previously developed a fluorescence-based, live-cell, high-content analysis (HCA) assay to examine drug efficacy against intracellular *M. tuberculosis* (15).

In this study, we screened 10,000 diverse small molecules by high throughput HCA and identified active compounds with low cytotoxicity. We selected a number of series for follow up based on their physicochemical properties and the novelty of the chemotypes and identified five chemical classes of interest. We tested available analogs for each series to complete a limited structure-activity relationship study. These new compound classes serve as starting points for the development of new series for antitubercular drug discovery.

## Methods

### Cell culture

Murine RAW 264.7 macrophages (ATCC TIB-71) were grown at 37°C in a humidified atmosphere containing 5% CO_2_ in RPMI 1640 medium supplemented with 5% fetal bovine serum, 1 mM sodium pyruvate solution, and 2 mM GlutaGro supplement (Corning). HepG2 cells (ATCC HB-8065) were cultured in Dulbecco-modified Eagle medium (DMEM), 10% fetal bovine serum (FBS), and 1× penicillin streptomycin solution (100 U/ml). *M. tuberculosis* H37Rv-LP (ATCC 25618) constitutively expressing codon-optimized *Ds*Red from plasmid pBlazeC8 (DREAM8) (16) was cultured at 37 °C in Middlebrook 7H9 medium containing 10% v/v OADC (oleic acid, albumin, dextrose, catalase) supplement (Becton Dickinson) and 0.05% w/v Tween 80 (7H9-Tw-OADC) plus 100 μg/mL hygromycin B.

### High content screening

Compounds were obtained in 384-well plates as 10 mM DMSO stock solutions and tested as described (15). Briefly, assay plates were prepared in clear bottom, black 384-well plates with 30 μL cRPMI and 0.6 μL compound in columns 3–22 (final concentration of 10 μM compound, 1% DMSO). 1% DMSO was used as the negative control, 10 mM isoniazid (INH) and 100 μM staurosporine (STA) were included as positive controls for anti-tubercular activity and cytotoxicity respectively. RAW267.4 cells were infected with *M. tuberculosis* DREAM8 as described (15) at an MOI of 1 for 24 h and extracellular bacteria removed by washing. Cells were recovered using Accumax, harvested, washed and resuspended in serum-free RPMI; 30 μL of infected cells were dispensed into each well at 3300 cells/well. Plates were incubated for 72 hours, 10 μL of 5X SYBR Green I was added and plates were imaged with an ImageXpress Micro High Content Screening System (Molecular Devices) using a 4x objective and FITC and Texas Red channels. MetaXpress was used to analyze images. The integrated intensity of *M. tuberculosis* or macrophages was calculated for each well. Growth inhibition was calculated for each test well by normalizing to the average integrated intensity of the DMSO control wells. IC_50_ was calculated as the compound concentration required to reduce bacterial growth by 50%. TC_50_ was calculated as the compound concentration required to reduce macrophage viability by 50%.

### Cytotoxicity

Cytotoxicity against HepG2 cells was measured after 72 hours. Cells were seeded in 384-well plates at 1800 cells per well. Compounds were added as a 10-point three-fold serial dilution after 24 h (final assay concentration of 1% DMSO). CellTiter-Glo® reagent (Promega) was added and relative luminescence units (RLU) measured. Data were normalized to the DMSO controls. Curves were fitted using the Levenberg–Marquardt algorithm; TC_50_ was calculated as the compound concentration required to reduce cell viability by 50%.

### Minimum inhibitory concentration

Minimum inhibitory concentrations (IC_90_) against *M. tuberculosis* were determined in liquid medium as described (17). Bacterial growth was measured after 5 days by OD_590_. IC_90_ was defined as the concentration of compound required to inhibit growth of *M. tuberculosis* by 90% and was determined using the Levenberg–Marquardt least-squares plot.

## RESULTS

### High throughput screening of a library of small molecules

We were interested in identifying new anti-tubercular molecules with activity against intracellular bacteria. We used our previously validated high throughput high content screen to test the MyriaScreen II library of 10,000 diverse molecules which is comprised of ∼60% singletons with diverse structural groups (TimTec/Sigma Aldrich). Molecules were tested at a single concentration of 10 μM (figure 1 A). We monitored cytotoxicity to the macrophages in the same wells to exclude false positives (since the bacteria are unable to replicate if the macrophages are dead). We identified 308 compounds with >70% bacterial growth inhibition; of these we discarded 134 with significant macrophage toxicity, defined as <70% macrophage survival (Figure 1B). This left 174 hit compounds that were non-cytotoxic but restricted the growth of *M. tuberculosis* in macrophages, a hit rate of 1.7%.

**Figure 1.**
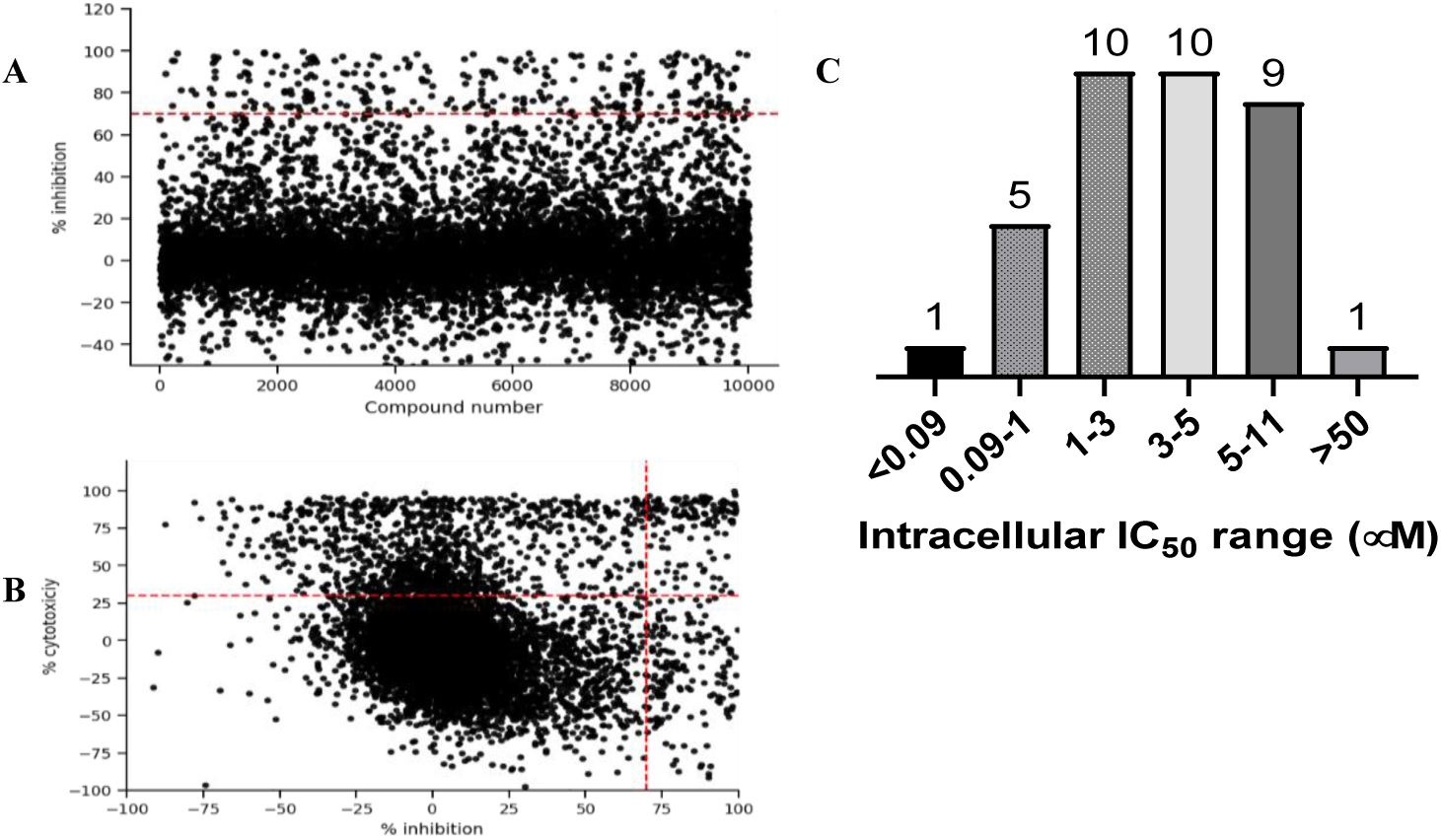
High content, high throughput screen against intracellular M. tuberculosis. (A). Compounds were tested for activity against RAW264.7 cells infected with *M. tuberculosis* at an MOI of 1. The primary screen was conducted using a single concentration of each molecule at 10 μM. Inhibition of bacterial growth and macrophage viability were determined for each compound after 72 hours. (A). Anti-tubercular activity measured as % inhibition of bacterial growth. (B) Cytotoxicity (% viability) was plotted compared to inhibition of bacterial growth (% Inhibition). Compounds in the top right quadrant were eliminated from the hit list due to cytotoxicity. (C) Selected hit compounds were purchased and tested in dose response to determine IC_50_ (concentration required to inhibit bacterial growth by 50%). Number of compounds n each group are noted.

### Hit confirmation

We analyzed the 174 resulting hits and prioritized hit compounds based on their physicochemical properties including molecular weight (MW), calculated partition coefficient (cLogP), novelty, structural diversity, chemical tractability, and commercial availability. We selected 43 compounds for further analysis (Figure 2); we purchased new batches of compounds and determined the activity using a dose response format (Figure 1C). Seven compounds exhibited cytotoxicity against eukaryotic cells that hampered our ability to determine their anti-bacterial activity and were excluded from further study (Table 1). The IC_50_ of the remaining 36 hits was determined (Figure 1C); of these all but one (35/36) had confirmed activity with IC_50_ <50 μM against intracellular *M. tuberculosis* (Table 1). The majority of the hits (26) had an IC_50_ <5 μM, and 6 compounds were potent with an IC_50_ <1 μM (Figure 1C). The majority of the compounds had no cytotoxicity (TC_50_ >50 μM); a few compounds were cytotoxic at higher concentrations, but retained good selectivity (Table 1).

**Table 1.** Activity of primary hits. NC – not calculated due to cytotoxicity. ND – not determined. cLogP was calculated using Collaborative Drug Discovery (https://www.collaborativedrug.com/).

| Molecule ID | MW | clogP | Intracellular activity |  |
| --- | --- | --- | --- | --- |
|  |  |  | Anti-tubercular activity IC<sub>50</sub> (μM) | Cytotoxicity TC<sub>50</sub> (μM) |
| TPN-0006204 | 311.34 | 3.24 | 11 ± 0.5 (n=2) | >100 (n=2) |
| TPN-0006205 | 322.22 | 3.97 | 1.4 ± 0.64 (n=4) | 57 ± 23 (n=3) |
| TPN-0006206 | 426.99 | 6.41 | NC | 57 ± 20 (n=3) |
| TPN-0006207 | 350.27 | 5.00 | 6.3 ± 0.3 (n=2) | >100 (n=2) |
| TPN-0006209 | 379.28 | 4.89 | NC | 68 ± 0.50 (n=2) |
| TPN-0006211 | 285.39 | 2.80 | 7.6 ± 3.5 (n=2) | 38 ± 6.0 (n=2) |
| TPN-0006216 | 330.20 | 4.55 | 2.8 ± 0.78 (n=4) | >100 (n=4) |
| TPN-0006217 | 276.35 | 3.32 | 4.9 ± 2.1 (n=4) | 75 ± 16 (n=4) |
| TPN-0073027 | 323.41 | 3.69 | 5.8 ± 2.1 (n=5) | >100 (n=5) |
| TPN-0073061 | 320.19 | 2.92 | 1.3 ± 0.15 (n=4) | 18 ± 4.3 (n=4) |
| TPN-0073062 | 272.37 | 4.21 | 0.39 ± 0.13 (n=4) | >100 (n=4) |
| TPN-0073063 | 311.40 | 4.58 | $0.25 \pm 0.23$ (n=6) | $47 \pm 28$ (n=6) |
| TPN-0073559 | 332.42 | 4.29 | NC | $50 \pm 24$ (n=4) |
| TPN-0073560 | 384.88 | 5.85 | NC | $23 \pm 4.7$ (n=4) |
| TPN-0073562 | 319.19 | 4.53 | $2.9 \pm 0.52$ (n=3) | $16 \pm 11$ (n=3) |
| TPN-0073563 | 329.20 | 4.09 | $4.7 \pm 1.1$ (n=5) | $28 \pm 7.0$ (n=5) |
| TPN-0073565 | 355.17 | 4.82 | $7.0 \pm 1.8$ (n=4) | >100 (n=4) |
| TPN-0073571 | 399.29 | 5.15 | NC | $15 \pm 5.0$ (n=2) |
| TPN-0073576 | 332.42 | 4.39 | $1.1 \pm 0.36$ (n=4) | >100 (n=4) |
| TPN-0073578 | 272.14 | 4.19 | $1.7 \pm 0.85$ (n=4) | $11 \pm 2.2$ (n=4) |
| TPN-0073584 | 478.58 | 2.61 | $5.1 \pm 1.7$ (n=3) | >100 (n=2) |
| TPN-0073585 | 424.94 | 6.81 | $0.4 \pm 0.11$ (n=3) | $9.4 \pm 1.9$ (n=3) |
| TPN-0073588 | 456.53 | 4.67 | $6.5 \pm 1.7$ (n=3) | $67 \pm 8.5$ (n=2) |
| TPN-0089695 | 356.80 | 3.76 | $4.7 \pm 1.2$ (n=2) | $45 \pm 17$ (n=2) |
| TPN-0089788 | 367.19 | 3.25 | $2.5 \pm 0.57$ (n=5) | >100 (n=5) |
| TPN-0095025 | 292.80 | 5.17 | $4.4 \pm 1.7$ (n=3) | $26 \pm 7.1$ (n=3) |
| TPN-0095026 | 387.67 | 5.21 | NC | $19 \pm 12$ (n=2) |
| TPN-0095027 | 295.34 | 1.45 | $0.10 \pm 0.010$ (n=4) | $72 \pm 16$ (n=4) |
| TPN-0095028 | 248.30 | 2.49 | $4.4 \pm 1.3$ (n=2) | $54 \pm 20$ (n=2) |
| TPN-0095029 | 472.93 | 5.54 | >50 (n=3) | >50 (n=2) |
| TPN-0095030 | 228.68 | 3.11 | $6.9 \pm 1.9$ (n=3) | >100 (n=2) |
| TPN-0095031 | 361.31 | 2.41 | $0.036 \pm 0.007$ (n=5) | $40 \pm 25$ (n=5) |
| TPN-0095032 | 335.25 | 6.51 | $3.5 \pm 0.83$ (n=3) | $9.9 \pm 0.98$ (n=3) |
| TPN-0095033 | 370.42 | 1.15 | $3.4 \pm 2.6$ (n=3) | >100 (n=3) |
| TPN-0095034 | 431.27 | 4.32 | $8.4 \pm 2.3$ (n=4) | >100 (n=4) |
| TPN-0095035 | 462.47 | 4.78 | $1.0 \pm 0.02$ (n=3) | >50 (n=3) |
| TPN-0095036 | 379.30 | 5.22 | $3.5 \pm 0.97$ (n=3) | $27 \pm 2.8$ (n=3) |
| TPN-0095037 | 357.88 | 4.20 | $2.0 \pm 0.53$ (n=3) | $54 \pm 26$ (n=3) |
| TPN-0095038 | 273.13 | 3.26 | $2.5 \pm 0$ (n=2) | $72 \pm 15$ (n=2) |
| TPN-0095039 | 363.22 | 4.68 | $2.1 \pm 0.74$ (n=3) | $24 \pm 6.1$ (n=3) |
| TPN-0095040 | 434.50 | 4.74 | NC | $31 \pm 18$ (n=2) |
| TPN-0095041 | 339.41 | 4.36 | $3.6 \pm 0.50$ (n=3) | $13 \pm 5.5$ (n=3) |
| TPN-0095042 | 343.86 | 3.80 | $3.7 \pm 0.62$ (n=3) | $21 \pm 9.3$ (n=3) |

**Figure 2.**
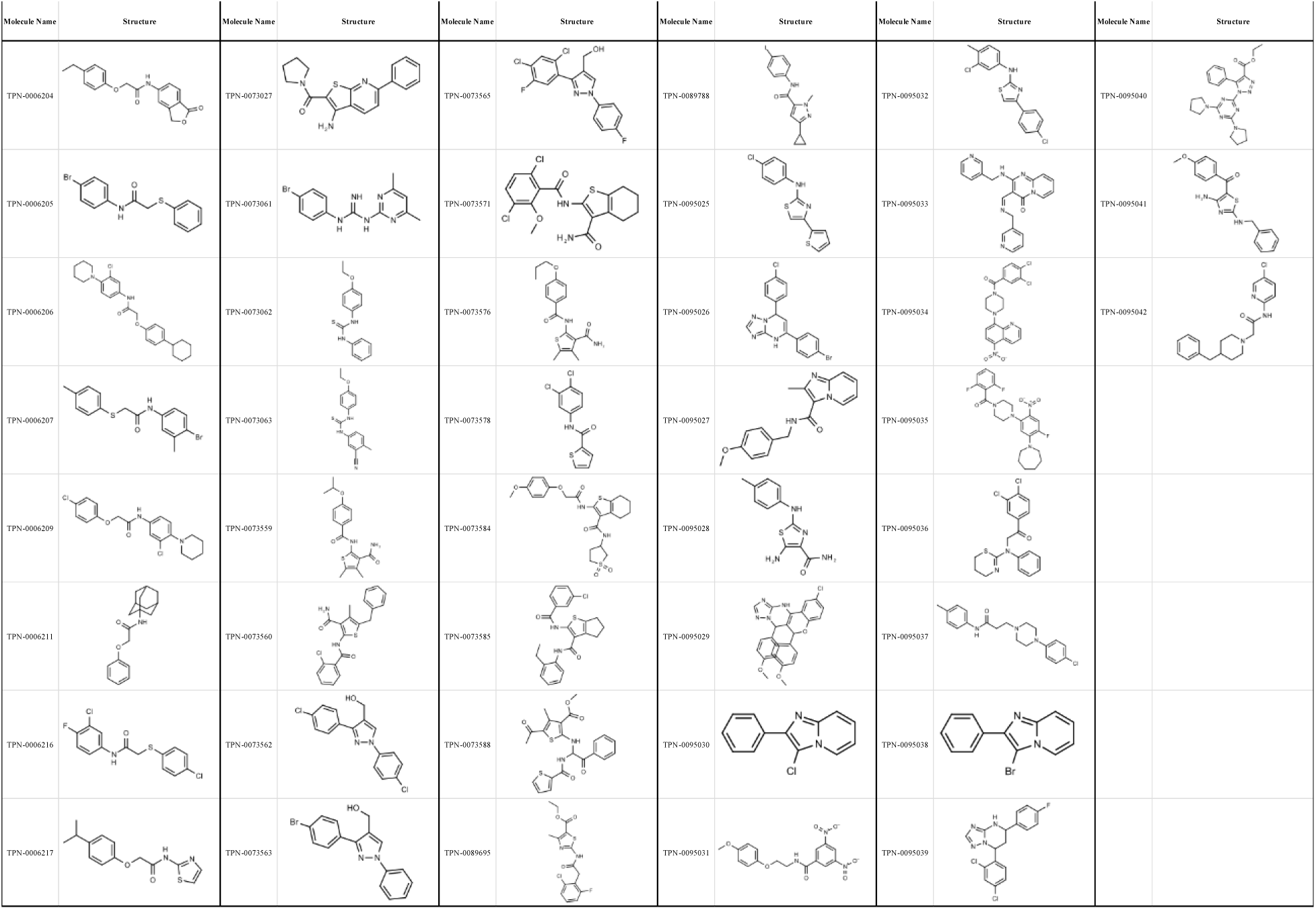
Structure of primary hits selected for follow up.

### Cytotoxicity

We further evaluated cytotoxicity using the HepG2 cell line cultured with either glucose or galactose in the medium (Table 2). HepG2 cells cultured with galactose rely on mitochondrial oxidative phosphorylation rather than glycolysis which increases their susceptibility to mitochondrial toxicants allowing us to identify compounds with mitochondrial toxicity (18). The majority of the compounds showed no difference in cytotoxicity between the two conditions with the exception of compound TPN-0073571 which had increased cytotoxicity against HepG2 cells in galactose (ratio of TC_50glu_/TC_50gal_ of ∼4) suggesting that this compound may impair mitochondrial function. Interestingly the compound showed similar cytotoxicity against the murine macrophages (TC_50_ = 15 ± 5 μM). Overall, cytotoxicity against the two cell lines (murine RAW and human HepG2 cells) was comparable (Table 2).

**Table 2.** Cytotoxicity and activity against extracellular *M. tuberculosis*.

| Molecule ID | HepG2 cytotoxicity $TC_{50}$ ($\mu\text{M}$) | | Anti-tubercular activity in axenic culture ($\mu\text{M}$) | |
| --- | --- | --- | --- | --- |
| | Glucose | Galactose | $IC_{50}$ | $IC_{90}$ |
| TPN-0006204 | $59 \pm 10$ (n=5) | $47 \pm 17$ (n=4) | $>20$ (n=2) | $>20$ (n=2) |
| TPN-0006205 | $>100$ (n=4) | $>100$ (n=3) | $>20$ (n=2) | $>20$ (n=2) |
| TPN-0006206 | $>100$ (n=2) | $>100$ (n=2) | $>20$ (n=2) | $>20$ (n=2) |
| TPN-0006207 | $>100$ (n=2) | $>100$ (n=2) | $>20$ (n=2) | $>20$ (n=2) |
| TPN-0006209 | $>100$ (n=2) | $>100$ (n=2) | $>20$ (n=2) | $>20$ (n=2) |
| TPN-0006211 | $53 \pm 1.5$ (n=2) | $43 \pm 1$ (n=2) | $>20$ (n=2) | $>20$ (n=2) |
| TPN-0006216 | $44 \pm 2$ (n=2) | $33 \pm 4$ (n=2) | $>20$ (n=2) | $>20$ (n=2) |
| TPN-0006217 | $91 \pm 3$ (n=2) | $64 \pm 1.5$ (n=2) | $>20$ (n=2) | $>20$ (n=2) |
| TPN-0073027 | $82 \pm 25$ (n=5) | $64 \pm 10$ (n=4) | $>20$ (n=3) | $>20$ (n=3) |
| TPN-0073061 | $22 \pm 4.0$ (n=3) | $22 \pm 3.1$ (n=3) | $6.3 \pm 0.15$ (n=2) | $8.2 \pm 0.1$ (n=2) |
| TPN-0073062 | $>100$ (n=4) | $>100$ (n=4) | $11 \pm 5.3$ (n=4) | $>20$ (n=4) |
| TPN-0073063 | 40 ± 11 (n=4) | 34 ± 6.5 (n=4) | 5 ± 1.5 (n=3) | 9.5 ± 3.2 (n=3) |
| TPN-0073559 | 49 ± 8.5 (n=3) | 41 ± 11 (n=3) | >20 (n=2) | >20 (n=2) |
| TPN-0073560 | >100 (n=2) | 42 ± 3.3 (n=3) | >20 (n=2) | >20 (n=2) |
| TPN-0073562 | 31 ± 2.1 (n=3) | 31 ± 1.9 (n=4) | 12 ± 0.5 (n=2) | 19 ± 1.5 (n=2) |
| TPN-0073563 | 34 ± 0 (n=3) | 34 ± 1.9 (n=3) | ND | ND |
| TPN-0073565 | 46 ± 2.8 (n=4) | 43 ± 2.1 (n=5) | >20 (n=2) | >20 (n=2) |
| TPN-0073571 | >100 (n=2) | 26 ± 5.0 (n=2) | ND | ND |
| TPN-0073576 | 83 ± 2.9 (n=3) | 92 ± 7 (n=3) | >20 (n=2) | >20 (n=2) |
| TPN-0073578 | 3.9 ± 1.1 (n=3) | 4.7 ± 0.54 (n=3) | >20 (n=2) | >20 (n=2) |
| TPN-0073584 | >100 (n=3) | >100 (n=3) | >20 (n=2) | >20 (n=2) |
| TPN-0073585 | >25 (n=3) | >25 (n=3) | >5 (n=2) | >5 (n=2) |
| TPN-0073588 | >100 (n=2) | >100 (n=3) | >20 (n=2) | >20 (n=2) |
| TPN-0089695 | >100 (n=5) | >100 (n=4) | 19 ± 0 (n=2) | >20 (n=2) |
| TPN-0089788 | 52 ± 18 (n=4) | 30 ± 12 (n=4) | >20 (n=2) | >20 (n=2) |
| TPN-0095025 | 35 ± 1.0 (n=2) | 28 ± 1.5 (n=2) | >20 (n=2) | >20 (n=2) |
| TPN-0095026 | 19 ± 3.5 (n=2) | 18 ± 5 (n=2) | 19 ± 1 (n=2) | >20 (n=2) |
| TPN-0095027 | 76 ± 12 (n=3) | 22 ± 2.2 (n=3) | 0.17 ± 0.02 (n=3) | 0.60 ± 0.20 (n=3) |
| TPN-0095028 | 54 ± 9.4 (n=3) | 62 ± 6.2 (n=3) | >20 (n=2) | >20 (n=2) |
| TPN-0095029 | >50 (n=2) | >50 (n=2) | >10 (n=2) | >10 (n=2) |
| TPN-0095030 | >100 (n=2) | >100 (n=2) | >20 (n=2) | >20 (n=2) |
| TPN-0095031 | 57 ± 6.2 (n=3) | 52 ± 0.47 (n=3) | 0.20 ± 0.07 (n=3) | 0.40 ± 0.14 (n=3) |
| TPN-0095032 | 23 ± 1.5 (n=2) | 19 ± 0.5 (n=2) | >20 (n=2) | >20 (n=2) |
| TPN-0095033 | >100 (n=3) | >100 (n=3) | >20 (n=1) | >20 (n=1) |
| TPN-0095034 | >100 (n=3) | >100 (n=3) | >20 (n=2) | >20 (n=2) |
| TPN-0095035 | 27 ± 0 (n=2) | 21 ± 2.0 (n=2) | 9.6 ± 0.1 (n=2) | >10 (n=2) |
| TPN-0095036 | 21 ± 3.4 (n=3) | 20 ± 0.47 (n=3) | >20 (n=1) | >20 (n=1) |
| TPN-0095037 | 84 ± 2.5 (n=2) | 56 ± 4 (n=2) | >20 (n=2) | >20 (n=2) |
| TPN-0095038 | >100 (n=2) | >100 (n=3) | 9.2 ± 1.0 (n=2) | >20 (n=2) |
| TPN-0095039 | 29 ± 7.7 (n=4) | 30 ± 3.2 (n=5) | 14 ± 2.1 (n=3) | 18 ± 1.4 (n=3) |
| TPN-0095040 | >50 (n=3) | >50 (n=3) | >10 (n=2) | >10 (n=2) |
| TPN-0095041 | 52 ± 5.5 (n=2) | 24 ± 0.5 (n=2) | 14 ± 1 (n=2) | >20 (n=2) |
| TPN-0095042 | 26 ± 3.0 (n=2) | 19 ± 1.0 (n=2) | >10 (n=2) | >10 (n=2) |

### Activity against *M. tuberculosis* in axenic culture

We wanted to determine if our compounds were selective for intracellular bacteria, so we determined the IC_90_ against *M. tuberculosis* in axenic culture. Only 12 compounds had IC_50_ > 20 μM against extracellular *M. tuberculosis*, with six of these having an IC_90_ of > 20 μM (Table 2). None of the compounds were more active against *M. tuberculosis* growing in axenic culture, with the extracellular/intracellular IC_50_ ratios ranging from 4 to 28.

### Identification of novel chemotypes

We performed structural clustering on the set of 174 hits in order to identify common and potentially privileged scaffolds. Compounds were grouped into series sharing the same scaffold/chemotype, each featuring 2 to 8 compounds with 8 singletons. Scaffolds used in the substructure searching, along with their frequency in the hit list and the entire library are depicted in Table 3.

**Table 3.**
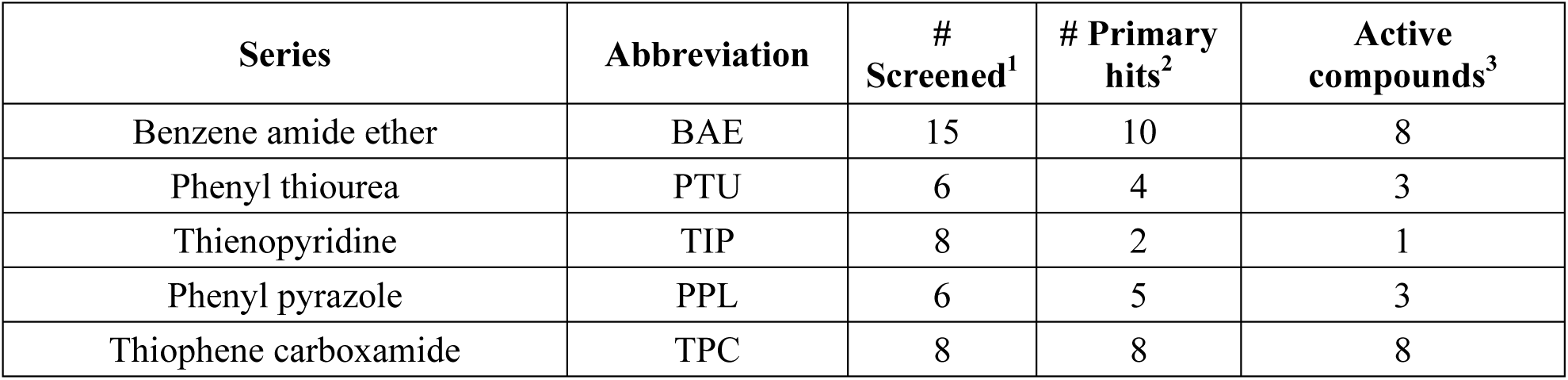
Series selected for follow-up. ^1^Number of compounds screened in the library. ^2^Number of compounds with activity in the single point screen. ^3^Number of confirmed hits (IC_50_ <20 μM).

The prioritization of compounds was not based solely on biological potency, but focus was placed on the quality/structural novelty of each chemotype, along with their physicochemical properties. We highlighted five chemotypes that were of interest for further development. A representative compound from each series is shown in Figure 3

**Figure 3.**
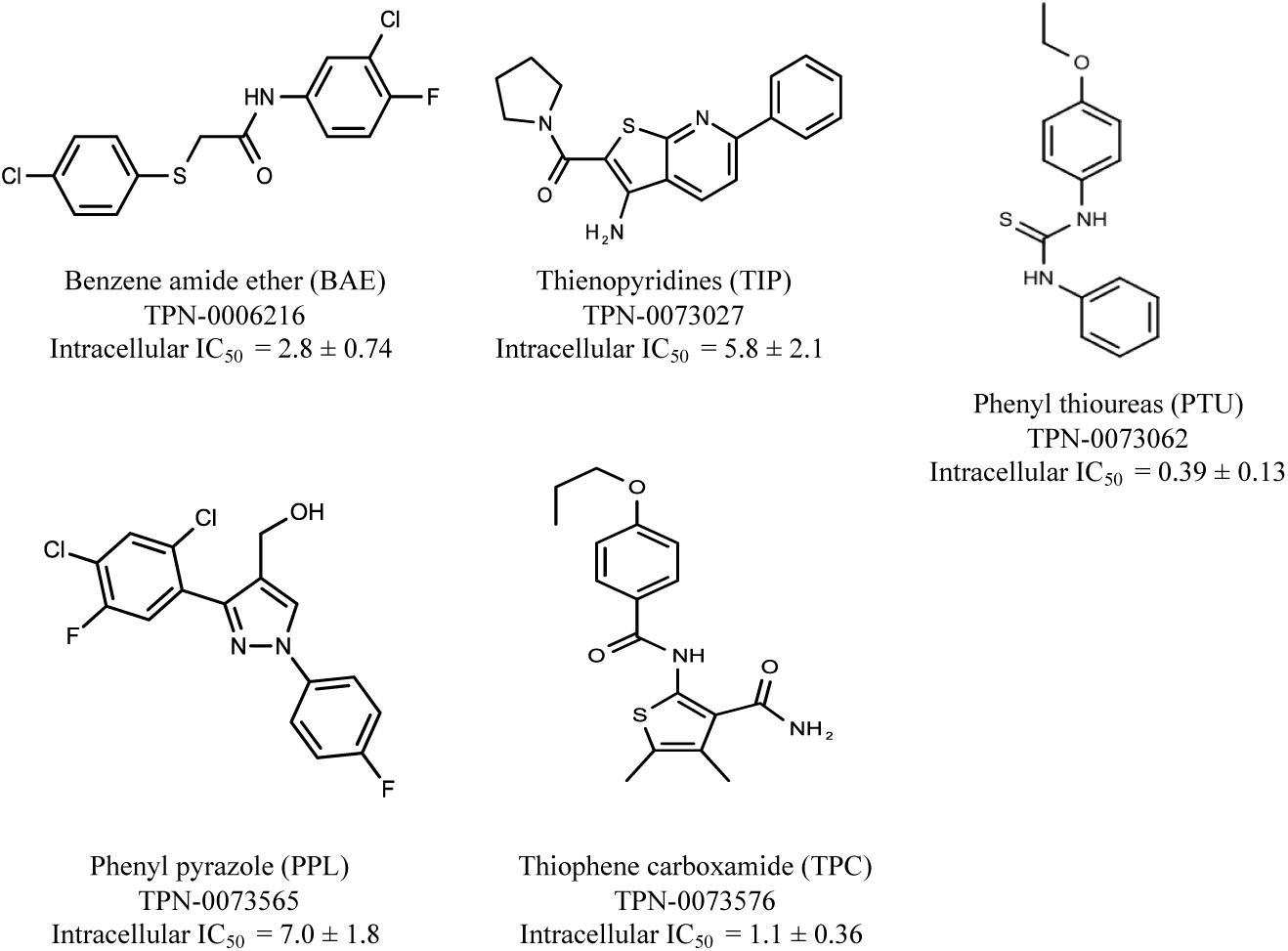
Representative molecules from selected series.

### Benzene-amide ether (BAE) series

We identified 15 compounds with a benzene amide ether scaffold in the primary screen, of which 10 were active (defined as >70% inhibition of bacterial replication) (Table 3). The BAE series has not previously been identified in phenotypic screens against replicating *M. tuberculosis*, although one compound from this series (**6**) had activity against non-replicating *M. tuberculosis* (19). In addition, compound **12** was identified as an inhibitor of *M. tuberculosis* with an IC_90_ of 6.9 μg/mL(20). We purchased 13 commercially-available compounds and determined intracellular activity against *M. tuberculosis*, although we did not see activity up to 20 μM. Eleven compounds had activity against intracellular *M. tuberculosis* (Table 4). Two compounds (**4** and **7**) were cytotoxic to macrophages and we could not calculate activity.

**Table 4.** Catalog structure-activity relationship of the BAE series. NC – not calculated due to cytotoxicity. cLogP was calculated using Collaborative Drug Discovery (https://www.collaborativedrug.com/).

| Benzene amide ether |  |  |  |  |  |  |  |  |  |
| --- | --- | --- | --- | --- | --- | --- | --- | --- | --- |
| $R_1-X-\text{CH}_2-\text{C}(=\text{O})-R_2$ | | | | | | | | | |
| Compound # | X | R<sub>1</sub> | R<sub>2</sub> | cLogP | Extracellular (μM) | Intracellular (μM) |  | HepG2 Cytotoxicity IC<sub>50</sub> (μM) |  |
|  |  |  |  |  | IC<sub>90</sub> | Mtb IC<sub>50</sub> | RAW TC<sub>50</sub> | Galactose | Glucose |
| 1 | S |  |  | 2.86 | >20 | 7.8 ± 1.6 | >100 | >100 | >100 |
|  |  |  |  |  | n=2 | n=2 | n=2 | n=2 | n=2 |
| 2 | O |  |  | 3.24 | >20 | 10 ± 0.5 | >100 | 47 ± 17 | 59 ± 10 |
|  |  |  |  |  | n=2 | n=3 | n=1 | n=4 | n=5 |
| 3 | S | Ph |  | 3.97 | >20 | 1.4 ± 0.6 | 57 ± 24 | >100 | >100 |
|  |  |  |  |  | n=2 | n=4 | n=3 | n=3 | n=4 |
| 4 | O |  |  | 6.41 | >20 | NC | 57 ± 20 | >100 | >100 |
|  |  |  |  |  | n=2 |  | n=3 | n=2 | n=2 |
| 5 | S |  |  | 5.00 | >20 | 6.3 ± 0.3 | >100 | >100 | >100 |
|  |  |  |  |  | n=2 | n=2 | n=2 | n=2 | n=2 |
| 6 | O |  |  | 3.59 | >20 | 13 ± 0 | 86 ± 1.0 | >100 | >100 |
|  |  |  |  |  | n=2 | n=2 | n=2 | n=2 | n=2 |
| 7 | O |  |  | 4.89 | >20 | NC | 68 ± 0.5 | >100 | >100 |
|  |  |  |  |  | n=2 |  | n=2 | n=2 | n=2 |
| 8 | O | Ph |  | 2.80 | >20 | 7.6 ± 3.5 | 38 ± 6.0 | 43 ± 1.0 | 53 ± 1.5 |
|  |  |  |  |  | n=2 | n=2 | n=2 | n=2 | n=2 |
| 9 | O |  |  | 2.50 | >20 | 8.2 ± 1.7 | >100 | 61 ± 4.0 | >100 |
|  |  |  |  |  | n=2 | n=2 | n=2 | n=2 | n=2 |
| 10 | O |  |  | 3.35 | >20 | 6.6 ± 1.6 | 75 ± 16 | 55 ± 5.0 | 93 ± 3.5 |
|  |  |  |  |  | n=2 | n=2 | n=2 | n=2 | n=2 |
| 11 | O |  |  | 3.58 | >20 | 12 ± 1.6 | 60 ± 27 | 57 ± 3.0 | 96 ± 2.5 |
|  |  |  |  |  | n=2 | n=2 | n=2 | n=2 | n=2 |
| 12 | S |  |  | 4.55 | >20 | 2.4 ± 0.5 | >100 | 33 ± 4.0 | 44 ± 2.0 |
|  |  |  |  |  | n=2 | n=2 | n=2 | n=2 | n=2 |
| 13 | O |  |  | 3.32 | >20 | 4.9 ± 2.1 | 70 ± 1.6 | 64 ± 1.5 | 91 ± 3.0 |
|  |  |  |  |  | n=2 | n=4 | n=4 | n=2 | n=2 |

Eight compounds showed good intracellular activity (IC_50_ <10 μM). There was a ten-fold difference between the most potent compound (**3**, IC_50_ = 1.4 μM) and the least potent compound **(13**, IC_50_ = 13 μM). Compounds generally showed low cytotoxicity against both RAW264.7 cells and HepG2 cells with either TC_50_ >100 μM or a selectivity index of >50 (TC_50_/IC_50_). Compound **3** and compound **12** were the most potent (IC_50_ = 1.4 μM and IC_50_ = 2.4 μM respectively). Although, compound **3** had some cytotoxicity, the selectivity index was 40. Compound **12** was not cytotoxic, with TC_50_ >100 μM and a selectivity index of >42. None of the compounds were active against replicating *M. tuberculosis* in axenic medium (IC_90_ >20 μM) demonstrating clear selectivity for intracellular bacteria.

We were able to determine some elements of the SAR for this series using the limited compound set available. Compounds with a sulfur atom as a linker were generally more potent than those with an oxygen as a linker, with a range of 1.4 -6.3 μM compared to 4.9-13 μM. The closest comparison was between **13** with the oxygen linker (IC_50_ = 4.9 μM) and compounds **3** and **12** (IC_50_ = 1.4 μM and 2.4 μM respectively). In addition, the two molecules with significant toxicity to macrophage cells contained the oxygen linker (**4** band **7**). Four compounds (**1, 3, 5** and **12)** with one or two substitution of halogen atoms on the amide phenyl ring (R_2_) exhibited good intracellular activity.

It seems that lipophilicity did not correlate with anti-tubercular activity. Although the most active compounds (**3** and **12**) had cLogP of 3.97 and 4.55 respectively, compound **13** with cLogP of 3.32 was only two-fold less active and compounds with cLogP <3 still retained good activity (IC_50_ <10 μM). An electron-withdrawing moiety on the para position of the amide phenyl group (R_2_) appears to be an essential requirement for good anti-mycobacterial activity. The amide phenyl ring (R_2_) is not critical to maintain antimycobacterial activity, since compounds **8** and **12** with saturated rings. Both compounds **8** and **10** with adamantane and piperidine groups respectively, had good activity (IC_50_ = 7.6 μM and IC_50_ = 6.6 μM). Further exploration of the length of the carbon chain or inclusion of other heteroatoms would expand the SAR, but such molecules are not available commerically and would require chemical synthesis.

### The phenylthiourea (PTU) series

We identified 6 compounds with a phenyl thiourea scaffold in the primary screen, of which 4 were active (Table 3). We purchased 5 commercially-available compounds and determined intracellular activity against *M. tuberculosis*. Four compounds had potent activity against intracellular *M. tuberculosis* (Table 5). One compound (**18**) was cytotoxic and we could not calculate activity. One compound (**17**) had significant cytotoxicity against macrophages (TC_50_ of 1.8 μM) and was also toxic for HepG2 cells (TC_50_ of 5.5 μM). The remaining three compounds had excellent potency (0.30 to 1.3 μM) and good selectivity. Compound **14** with a guanidine functionality as the core had higher cytotoxicity (SI = 14) than the two potent molecules (**15** and **16**) with the thiourea core (SI = >250 and >150 respectively).

**Table 5.**
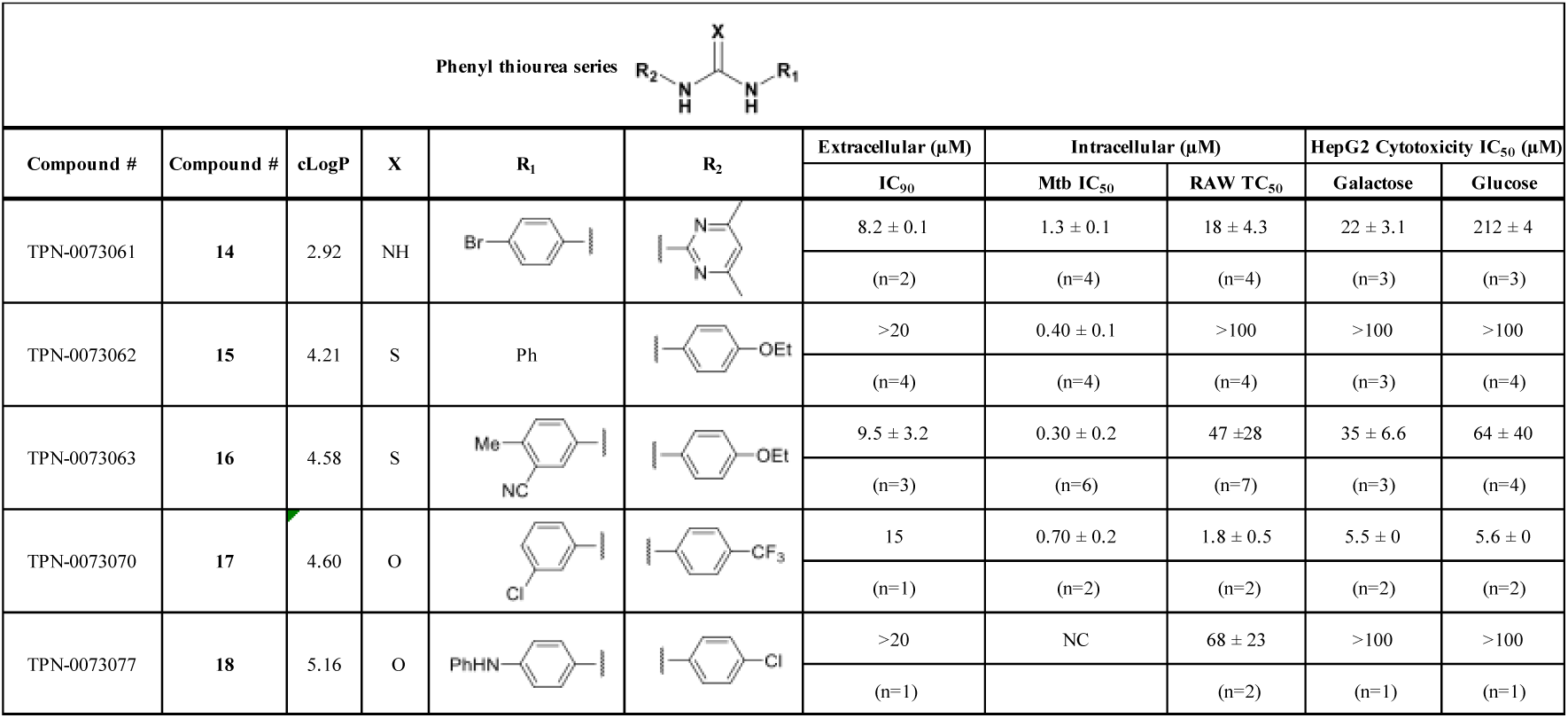
Catalog structure-activity relationship of the PTU series. NC – not calculated due to cytotoxicity. cLogP was calculated using Collaborative Drug Discovery (https://www.collaborativedrug.com/).

| Phenyl thiourea series |  |  |  |  |  |  |  |  |  |  |
| --- | --- | --- | --- | --- | --- | --- | --- | --- | --- | --- |
| Compound # | Compound # | cLogP | X | R<sub>1</sub> | R<sub>2</sub> | Extracellular (μM) | Intracellular (μM) |  | HepG2 Cytotoxicity IC<sub>50</sub> (μM) |  |
|  |  |  |  |  |  | IC<sub>90</sub> | Mtb IC<sub>50</sub> | RAW TC<sub>50</sub> | Galactose | Glucose |
| TPN-0073061 | **14** | 2.92 | NH |  |  | 8.2 ± 0.1 | 1.3 ± 0.1 | 18 ± 4.3 | 22 ± 3.1 | 212 ± 4 |
|  |  |  |  |  |  | (n=2) | (n=4) | (n=4) | (n=3) | (n=3) |
| TPN-0073062 | **15** | 4.21 | S | Ph |  | >20 | 0.40 ± 0.1 | >100 | >100 | >100 |
|  |  |  |  |  |  | (n=4) | (n=4) | (n=4) | (n=3) | (n=4) |
| TPN-0073063 | **16** | 4.58 | S |  |  | 9.5 ± 3.2 | 0.30 ± 0.2 | 47 ± 28 | 35 ± 6.6 | 64 ± 40 |
|  |  |  |  |  |  | (n=3) | (n=6) | (n=7) | (n=3) | (n=4) |
| TPN-0073070 | **17** | 4.60 | O |  |  | 15 | 0.70 ± 0.2 | 1.8 ± 0.5 | 5.5 ± 0 | 5.6 ± 0 |
|  |  |  |  |  |  | (n=1) | (n=2) | (n=2) | (n=2) | (n=2) |
| TPN-0073077 | **18** | 5.16 | O |  |  | >20 | NC | 68 ± 23 | >100 | >100 |
|  |  |  |  |  |  | (n=1) |  | (n=2) | (n=1) | (n=1) |

In contrast to the BAE series, the PTU molecules were also active against extracellular bacteria; 3 of the 5 moelcules had IC_90_ <20 μM. In general, PTUs were 5-10-fold more active against intracellular bacteria. To the best of our knowledge, this is the first report of these compounds as antitubercular agents although other thiourea derivatives have antitubercular activity(21–24).

### The phenyl pyrazole (PPL) series

We identified 6 compounds with a phenyl pyrazole scaffold in the primary screen, of which 5 were active (Table 3). We purchased 4 commercially-available compounds and determined intracellular activity against *M. tuberculosis*. All four compounds (**19-22**) had activity against intracellular *M. tuberculosis* (Table 6). Although the compounds demonstrated some cytotoxicity against HepG2 cells and/or macrophages, the selectivity index was good (>5). These sets of compounds have 1,3,4-trisubstituted pyrazoles where C-4 bears a hydroxymethyl group. Compound **19** was the most potent with good intracellular activity (IC_50_ = 2.9 μM) and > 30-fold selectivity against HepG2 cells. Compound **21** also showed weak activity (IC_50_ = 18.5 μM) against axenically-grown *M. tuberculosis*. Previous work has identified similar pyrazole-containing compounds with anti-tubercular activity that target MmpL3. In these studies, a 1,3,5-trisubstituted pyrazole as a central core had good activity against axenically-cultured *M. tuberculosis* (25). Our moelcules differ in that they only have weak activity against extracellular bacteria. The molecules we identify here are phenyl pyrazoles which have not been reported previously to have activity against either intracellular or extracellular bacteria.

**Table 6.**
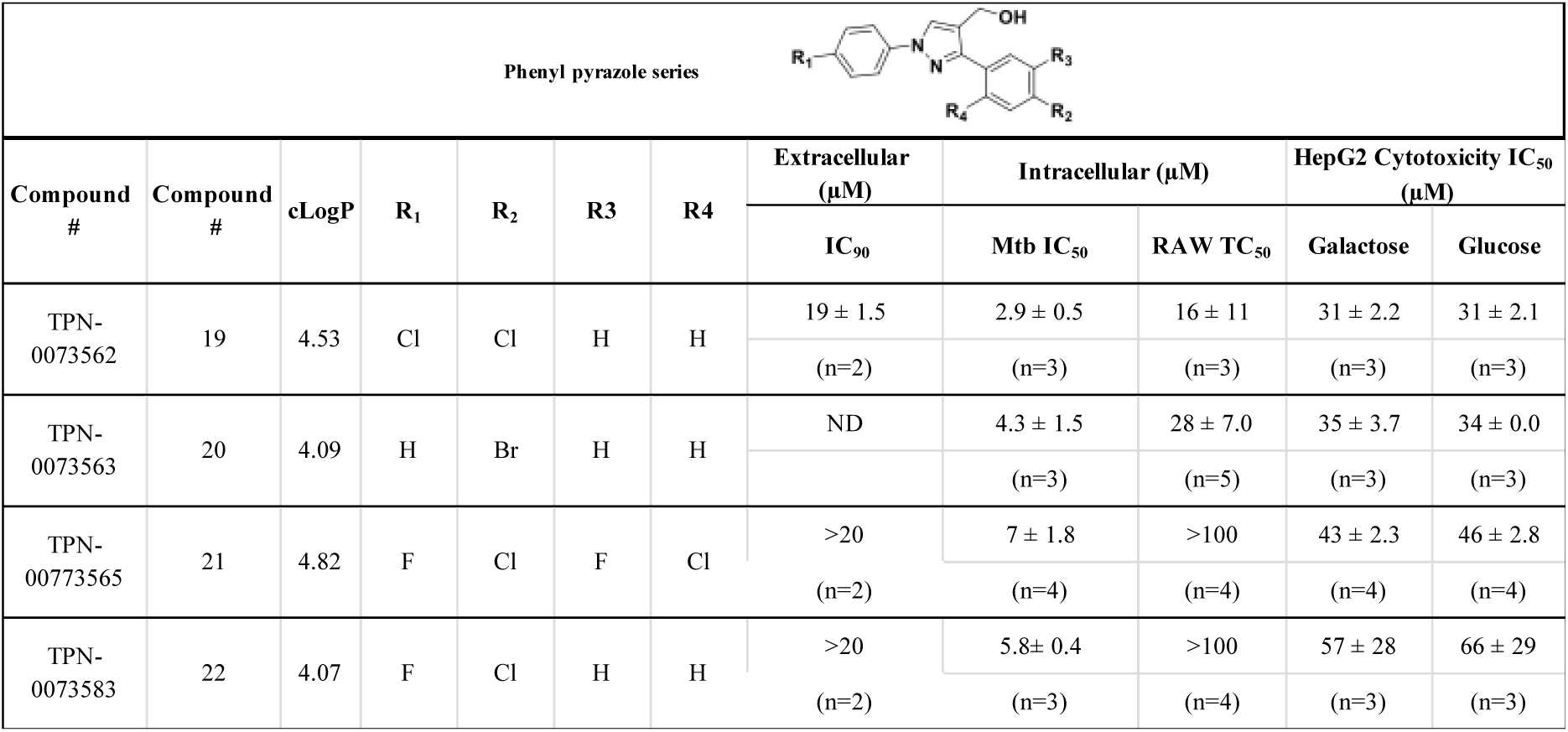
Catalog structure-activity relationship of the PPL series. ND – not determined. cLogP was calculated using Collaborative Drug Discovery (https://www.collaborativedrug.com/).

| Phenyl pyrazole series |  |  |  |  |  |  |  |  |  |  |  |
| --- | --- | --- | --- | --- | --- | --- | --- | --- | --- | --- | --- |
| Compound # | Compound # | cLogP | R<sub>1</sub> | R<sub>2</sub> | R<sub>3</sub> | R<sub>4</sub> | Extracellular (μM) | Intracellular (μM) |  | HepG2 Cytotoxicity IC<sub>50</sub> (μM) |  |
|  |  |  |  |  |  |  | IC<sub>90</sub> | Mtb IC<sub>50</sub> | RAW TC<sub>50</sub> | Galactose | Glucose |
| TPN-0073562 | 19 | 4.53 | Cl | Cl | H | H | 19 ± 1.5 | 2.9 ± 0.5 | 16 ± 11 | 31 ± 2.2 | 31 ± 2.1 |
|  |  |  |  |  |  |  | (n=2) | (n=3) | (n=3) | (n=3) | (n=3) |
| TPN-0073563 | 20 | 4.09 | H | Br | H | H | ND | 4.3 ± 1.5 | 28 ± 7.0 | 35 ± 3.7 | 34 ± 0.0 |
|  |  |  |  |  |  |  |  | (n=3) | (n=5) | (n=3) | (n=3) |
| TPN-00773565 | 21 | 4.82 | F | Cl | F | Cl | >20 | 7 ± 1.8 | >100 | 43 ± 2.3 | 46 ± 2.8 |
|  |  |  |  |  |  |  | (n=2) | (n=4) | (n=4) | (n=4) | (n=4) |
| TPN-0073583 | 22 | 4.07 | F | Cl | H | H | >20 | 5.8 ± 0.4 | >100 | 57 ± 28 | 66 ± 29 |
|  |  |  |  |  |  |  | (n=2) | (n=3) | (n=4) | (n=3) | (n=3) |

### The thiophene carboxamide series (TPC)

We identified 8 compounds with a thiophene carboxamide scaffold in the primary screen, of which all 8 were active (Table 3). We purchased 8 commercially-available compounds (**23-30**) and determined intracellular activity against *M. tuberculosis*. Five compounds had activity against intracellular *M. tuberculosis* with good acitivity of IC_50_ <10 μM (Table 7). Three compounds (**23-25**) were cyotoxic and we could not calculate activity.

**Table 7.**
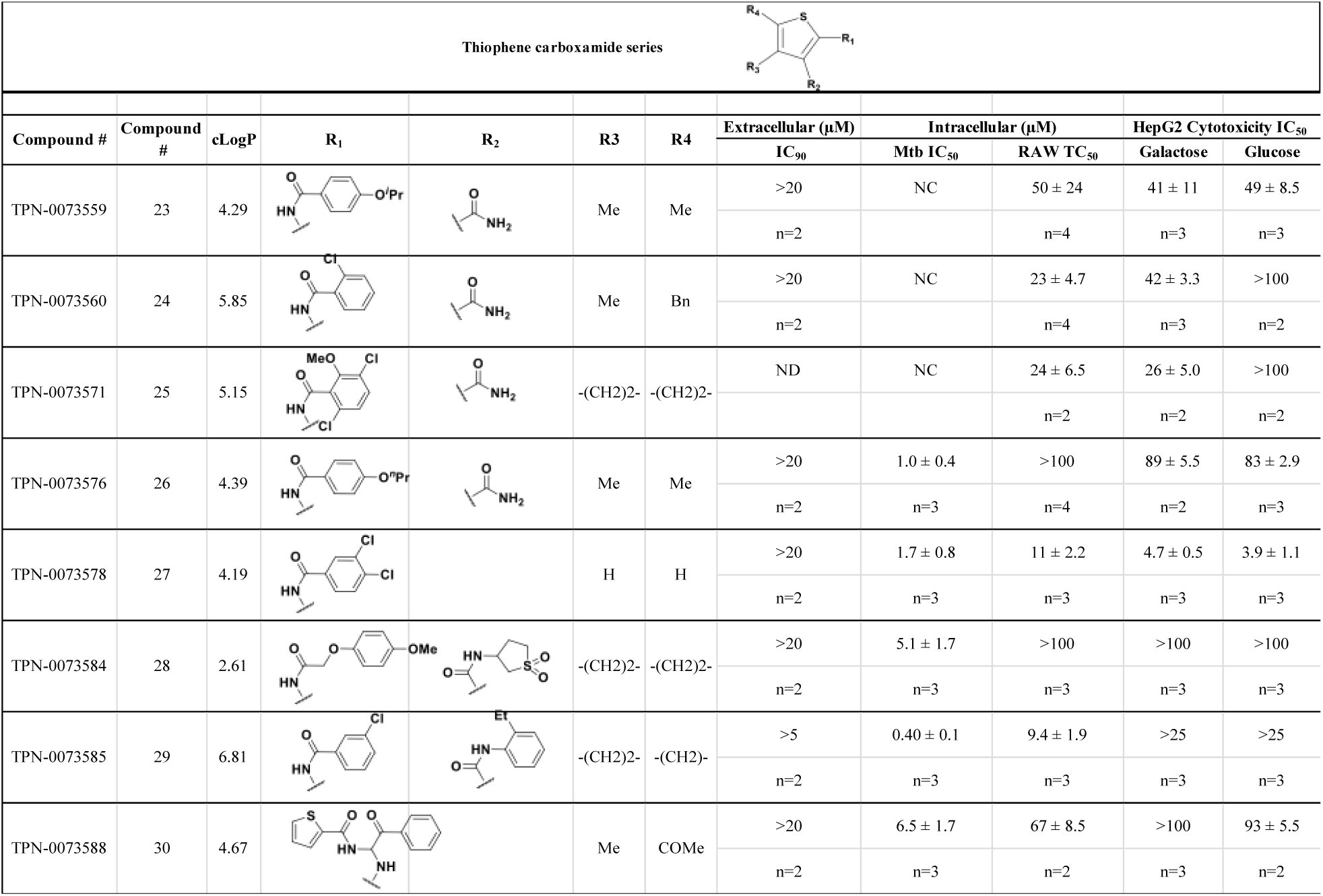
Catalog structure-activity relationship of the TPC series. ND – not determined. NC – not calculated due to cytotoxicity. cLogP was calculated using Collaborative Drug Discovery (https://www.collaborativedrug.com/).

Three compounds (**26, 27** and **29**) showed good intracellular potency with IC_50_ <5 μM. All except **26** and **28** were cytotoxic against macropahges with IC_50_ ranging from 9-67 μM; compound **27** was the most toxic and also showed cytotoxicity against HepG2 cells (IC_50_ = 4.7 μM). However, despite this, compounds **26, 28, 29**, and **30** all had good selectivity (SI>10) – values for HepG2 were >80, 20, >60 and 15 respectively. Two compounds (**24** and **25**) were more cytotoxic against HepG2 cells in galactose media (ratio ∼4) suggesting that they may impair mitochondrial function. However, these were also toxic against macrophages and were not considered active. None of the compounds had any activity against extracellular *M. tuberculosis*. To the best of our knowledge, this is the first report of these compounds as antitubercular agents.

### The thienopyridines series (TIP)

We identified 8 compounds with a thiophene carboxamide scaffold in the primary screen, of which 2 were active (Table 3).

We purchased 7 commercially-available compounds (**31-37**) and determined intracellular activity against *M. tuberculosis* Analogs varied in terms of substitution on the pyridine ring but all possessed the amino group at the C-3 position and a carboxamide substitution on the thiophene ring. Four compounds had activity against intracellular *M. tuberculosis* (Table 8), but only two had activity <10 μM (compounds **34** and **36)**. Compound **34** bearing a ketone moiety had weak activity. Two compounds (**31** and **35**) were not active. Compounds were selective, but had no activity against axenically cultured bacteria.

**Table 8.**
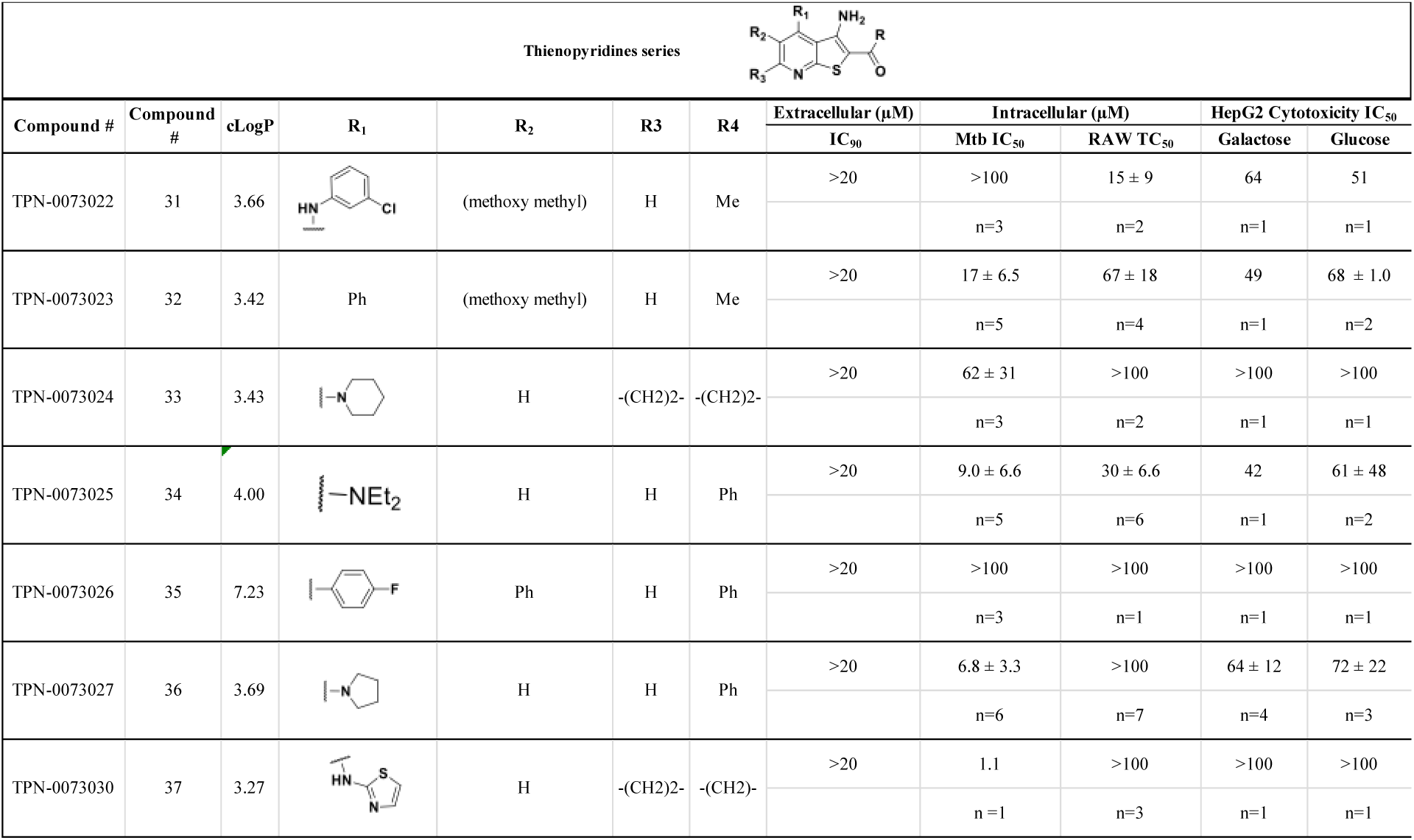
Catalog structure-ctivity relationship of the TIP series. cLogP was calculated using Collaborative Drug Discovery (https://www.collaborativedrug.com/).

## DISCUSSION

We screened a diversity in order to find novel chemical matter that targets intracellular *M. tuberculosis*. We identified a large number of hits with a hit rate of 1.7%. This is comparable to the hit rate we see with other whole cell screens and in other groups. We were able to follow up on several series to conduct limited catalog SAR as a first evaluation of tractability. The majority of the hit series we followed up from in this screen were more active against intracellular bacteria and in some cases lacked any extracellular activity. Since we focused on identifying new scaffolds, this was largely expected, since scaffolds with known anti-tubercular activity from previous phenotypic screens would have been deprioritized in our selection process. For the five series we prioritized, this is the first report of antimycobacterial activity of the majority of these compounds. This highlights the power of the screen assay in identifying novel compounds that are conditionally active against intracellular *M. tuberculosis*. Compounds that are only active against intracellular bacteria could be targeting pathways that are only essential for growth within the macrophages. Alternatively, they could target host pathways that enhance bacterial clearance or could be accumulated and/or metabolized within the macrophage. Further work to identify targets and mode of action is warranted on series with attractive properties.

Early identification of potentially toxic compounds during library screening has a significant impact on both the cost and the success rate of the drug discovery process. We were able to identify cytotoxic compounds rapidly and to exclude componuds that may look active due to killing the macrophages rather than anti-bacterial activity.. Out of the 308 hits that we identified as mycobacterial growth inhibitors, we excluded 134 molecules with significant toxicity to the macrophages representing almost a third of the hits. We also saw that compounds had comparable cytotoxicity against both macrophage and HepG2 cell lines, demonstrating the power of this assay in predicting potential cytotoxicity of the compounds early in the screening process.

## CONCLUSION

We have identified five novel and tractable chemical scaffolds with promising activity against intracellular *M. tuberculosis* using a high throughput, high content screen. The mechanism of action and targets remains to be characterized for each series. However, these series represent attractive starting points for lead generation and full structure-activity relationship studies.

## ACKNOWLEDGEMENTS

We thank Yulia Ovechkina and Sultan Chowdhury for technical assistance and useful discussion.

Research reported in this publication was supported by NIAID of the National Institutes of Health under award number R01AI132634. The content is solely the responsibility of the authors and does not necessarily represent the official views of the National Institutes of Health.

